# NORTH: a highly accurate and scalable Naive Bayes based ORTHologous gene cluster prediction algorithm

**DOI:** 10.1101/528323

**Authors:** Nabil Ibtehaz, Shafayat Ahmed, Bishwajit Saha, M. Sohel Rahman, Md Shamsuzzoha Bayzid

## Abstract

**Background:** Identifying orthologous genes plays a pivotal role in comparative genomics as the orthologous genes remain less diverged in the course of evolution. However, identifying orthologous genes is often difficult, slow, and idiosyncratic, especially in the presence of multiplicity of domains in proteins, evolutionary dynamics, multiple paralogous genes, incomplete genome data, and for distantly related species.

**Results:** We present NORTH, a novel, automated, highly accurate and scalable machine learning based orhtologous gene cluster prediction method. We have utilized the biological basis of orthologous genes and made an effort to incorporate appropriate ideas from machine learning (ML) and natural language processing (NLP). NORTH outperforms the frequently used existing orthologous clustering algorithms on the OrthoBench benchmark, not only just quantitatively with a high margin, but qualitatively under the challenging scenarios as well. Furthermore, we studied 12,55,877 genes in the largest 250 orthologous clusters from the KEGG database, across 3,880 organisms comprising the six major groups of life. NORTH is able to cluster them with 98.48% precision, 98.43% recall and 98.44% *F*_1_ score.

**Conclusions:** This is the first study that maps the orthology identification to the *text classification* problem, and achieves remarkable accuracy and scalability. NORTH thus advances the state-of-the-art in orthologous gene prediction, and has the potential to be considered as an alternative to the existing phylogenetic tree and BLAST based methods.

## 1 Introduction

The concept of homology has been adopted as the basis of phylogenetics and comparative genomics. Despite many arguments about its interpretation [1], homology can be defined as a similarity relationship between features that is due to shared ancestry. Homologous sequences can be further divided into orthologs and paralogs, according to how they diverged from their common ancestor. As originally defined in [2], homologous sequences that are derived through speciation are called orthologs. On the contrary, paralogs are homologous sequences that are derived from a duplication event [2, 3]. Detection of orthologs is of utmost importance in many fields of biology, especially in the annotation of genomes, functional genomics, evolutionary and comparative genomics. The pattern of genetic divergence can be used to trace the relatedness of organisms. Since orthologs are originated from speciation, they tend to preserve similar molecular and biological functions [4]. Paralogs, however, are originated from gene duplications and tend to deviate from their ancestral behaviour and functions [5, 6]. Nevertheless, orthologs are not necessarily identical in nature, as some orthologous genes can substantially diverge even among closely related organisms [7]. The converse argument is also valid – identically functioning genes are not necessarily orthologs [8, 9].

To date, considerable attentions have been drawn towards developing methods to identify orthologs and paralogs and experimented using complex datasets [10, 11, 12, 13, 14, 15]. These methods can be primarily categorized into phylogenetic tree based approaches and sequence similarity based methods (also known as “graph-based” methods). Tree-based methods are the classical approach for orthology inference that seek to reconcile gene and species trees. In most cases gene and species trees have different topologies due to evolutionary events acting specifically on genes, such as duplications, losses, lateral transfers, or incomplete lineage sorting [16]. Goodman et al. [17] resolved these incongruences by explaining them in terms of speciation, duplication, and loss events on gene trees with respect to species trees. Therefore, orthology/paralogy inference can be reduced to gene tree and species tree reconciliation problem. Sequence based approaches, on the other hand, exploit the relative similarity between the orthologous genes. These methods are primarily based on BLAST search for finding the pairs having the highest sequence identity, as in Inparanoid [18], OrthoMCL [19], BBH [20], RSD [21], OrthoFinder [22], morFeus [23], Ortholog-Finder [24], OrthoVenn [25], PorthoMCL [26] etc. They are also known as “graph-based” techniques as they create a graph with genes as the vertices and pairwise scores as the edge weights and use graph based clustering techniques for finding the orthologs. Most of these methods also include a post-processing step involving various heuristics.

In recent years, a number of machine learning based algorithms have been proposed for the inference of orthologous relationships among genes. However, all of those studies were constrained to a limited number of organisms. For example, Galpart *et al.* [27] studied three pairs of related yeast species, Sutphin and Mahoney [28] considered six eukaryotic species and Towfic *et al.* [29] analyzed just four organisms. Not only the datasets were very small, these approaches are not interpretable as a parallel to the biological insights.

These three types of methods circumvent some of the challenging issues in orthology inference, but have limitations of their own. One of the major issues with the existing methods is that they are prone to *precision-recall trade-off* (i.e., they fail to simultaneously achieve high precision and recall) [11, 12, 30]. Existing methods are computationally too demanding to analyze the ever-increasing number of genomes and thus often require access to supercomputers [15, 31]. Some of the existing methods are difficult to apply on eukaryotes [19], while some are restricted only to pairwise orthology detection [20, 27]. Tree-based methods suffer less from differential gene-loss and varying rates of evolution than BBH methods [32, 33], but genome-wide reconstruction of gene trees and species trees are computationally very demanding. Moreover, this approach is very sensitive to gene tree estimation error [34], and often performs worse than the sequence similarity based methods, and requires manual curation [11, 35]. Sequence similarity based methods can overcome many of these issues and can perform well on closely related organisms [36]. However, very often they do not take the evolutionary ortholog divergence into consideration. Due to this lack of evolutionary information, they tend to mistakenly detect homoplasious paralogs as orthologs [37]. These methods may achieve relatively high precision, albeit at the cost of a relatively low recall [20, 38, 39]. Moreover, BBH based methods can only account for one-to-one orthologs and may potentially miss true orthologs where one-to-many and many-to-many relationships are required to properly describe the orthology relationships [15, 36]. On the other hand, machine learning techniques, despite having an astounding potential, have only been validated on a small numbers of species.

In this study, we propose a novel method for predicting orthologous gene clusters which potentially overcomes most of the challenging issues and idiosyncrasies of the existing techniques. Taking inspirations from the widely used BLAST search, we treat this as a “text classification problem” and show that a Multinomial Naive Bayes algorithm with a bag-of-words model is able to predict orthology relationships to a near perfect accuracy even when we are dealing with extremely unbalanced data with more than a million of genes on a diverse as well as distant set of organisms.

## 2 Materials and Methods

### 2.1 Rationale: from BLAST search to text classification

Sequence-based methods rely on the assumption that the protein sequences obtained from orthologous genes tend to be quite similar since they are more conserved in the course of evolution [40]. These pipelines revolve mostly around a BLAST search [41], and often some parameter tuning and post-processing are performed for further refinement [42].

BLAST (Basic Local Alignment Search Tool) divides the entire sequence into small words or *k*-mers, and searches the database for finding sequences that resemble the query sequence in terms of the *k*-mer distribution [43]. After filtering out low complexity regions, BLAST constructs *k*-mers and scores them using a substitution matrix, which later seeds a likely un-gapped alignment. The seed alignments are extended to the left and to the right until the accumulated total score declines below a threshold, and the resultant alignment is computed thereby.

Quite interestingly, the popular “Text Classification Problem” [44] in NLP apparently resembles the BLAST protocol. Here, a text document is tokenized into words [45] and based on the word frequency the most similar category is selected, preferably with a classifier. Thus, a parallel can be drawn between the two by treating *k*-mers as words. This has been the initial enthusiasm for this work – replacing computationally demanding BLAST search procedure with a standard text classification protocol, coalescing the biological knowledge of orthology inference with the concepts in NLP.

### 2.2 NORTH: Ortholog cluster prediction through text classification

NORTH takes a protein sequence as input, and puts it in one of the predefined ortholog clusters using the following steps (please see Fig. 1 for an overview of the pipeline).

**Figure 1:**
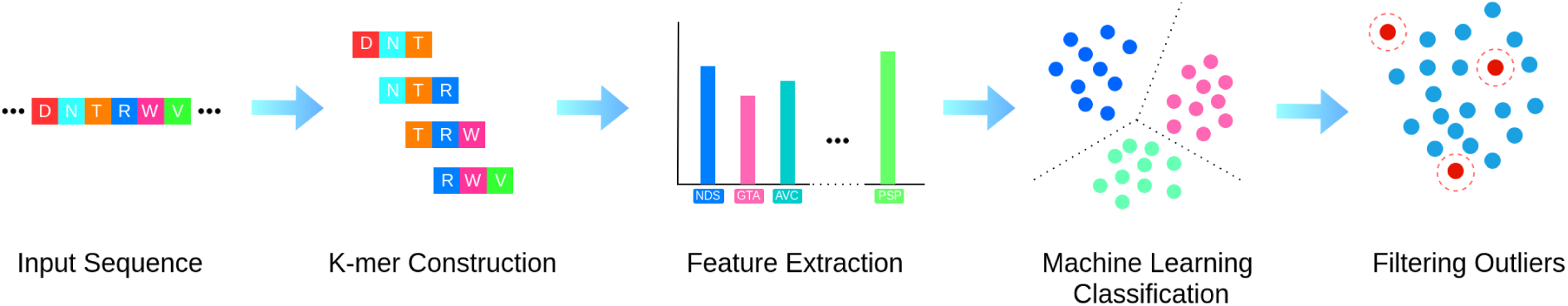
Algorithmic pipeline of NORTH. NORTH breaks the input protein sequence of the gene into *k*-mers, and uses the *k*-mer frequencies as the features to the multinomial Naive Bayes classifier to classify the input sequence into one of the predefined orthologous clusters. Consequently, it reports whether the gene is a member of the predefined orthologous clusters or an outlier.

#### 2.2.1 *k*-mer construction

We break the input sequence into *k*-mers. Although the common practice is to take *k* = 3 for doing a BLAST on amino acid sequence, *k* is a tunable parameter in our algorithm. The intuition is that higher values of *k* will enable the algorithm to capture more context, which will aid in considering diversified data, but may cause overfitting, compelling the algorithm to focus on some specific patterns. Conversely, a low *k* will dissipate coarse details, making the method more relaxed.

#### 2.2.2 Feature extraction

After breaking down the input protein sequence into *k*-mers, we compute the frequency distribution of the diversified *k*-mers. We use the *k*-mer frequencies as features for our algorithm, which is quite similar to the very popular bag-of-words model [46] widely used in natural language processing (NLP). NORTH does not resort to any other sophisticated feature engineering techniques.

#### 2.2.3 Machine learning classification

Every gene in an ortholog cluster is mutually orthologous to each other. Thus our objective is to map each sequence into one of the predefined ortholog clusters. As part of our attempt to coalesce the approaches of orthologous gene clustering and text classification, we use a multinomial Naive Bayes classifier [47] to cluster the genes based on the computed features.

#### 2.2.4 Filtering outliers

A supervised classifier is expected to infer a predefined number of orthologous clusters, leading to an erroneous decision for a gene outside these clusters. Hence, in order to assert the reliability of the proposed algorithm, it is required to identify potential outlier genes, i.e., genes that are outside of the clusters under consideration.

Analysis of the pattern of the distribution of cluster probabilities obtained from the Naive Bayes algorithm reveals that in case of a valid clustering, the probabilities follow a sharp, needle in a haystack pattern. On the contrary, while predicting the cluster of an outlier gene, the probability distribution is comparatively flat and scattered. Therefore, comparing the pattern of cluster probabilities becomes a viable option for NORTH to identify outlier genes. In order to formulate the pattern of the cluster probabilities, we normalize all probability values to the range [0, 1]. Then, various statistical attributes of the distribution, namely, mean, standard deviation, sum, skewness and kurtosis are extracted [48]. Using these as features, we train a Random Forest classifier, which is an ensemble of decision trees [49, 50], to infer whether a prediction is valid or represents an outlier gene.

### 2.3 Scalability

One of the notable attributes of the popular BLAST search is that it scales with the number of CPU cores [51]. As a result, to present NORTH as an alternative to BLAST-based approaches, we propose a scalable implementation of NORTH, which will aid clustering of plethora of genes. The computation of NORTH is divided automatically into a number of independent jobs, which are computationally less expensive and can be performed in parallel. After the computations are finished, they are merged together to generate the final result. In a single computer environment, these jobs can be performed sequentially, or in parallel on different threads or processes when considering a multi-core architecture. However, such a map-reduce like framework will excel the most in a distributed computing environment. More details on our scalable implementation can be found in Appendix A.

### 2.4 Selection of k

As we increase the values of *k*, analyzing the data becomes computationally too expensive since we have to consider around 20^*k*^ *k*-mers. Although scalable NORTH is able to handle it, the data preprocessing became exponentially complicated. Thus, we kept our experiments limited up to *k* ≤ 6, which were performed on the data from the KEGG database. In most of the cases, the best results are obtained for *k* = 5 as illustrated in Fig. 4. The only exception is the presence of prokaryotic genes, where *k* = 6 achieves the better results. However, *k* = 5 still holds up quite well in those cases, with a slight fall of accuracy (less than 0.5%). Thus, NORTH is robust and not much sensitive to the values of *k*, and considering the trade-off between computational requirement and accuracy, *k* = 5 appears to be a suitable option. It is worth noting that results from experiments with *k* = 7, is not significantly different from that for *k* = 6 (see supplementary material SM.xls).

**Figure 2:**
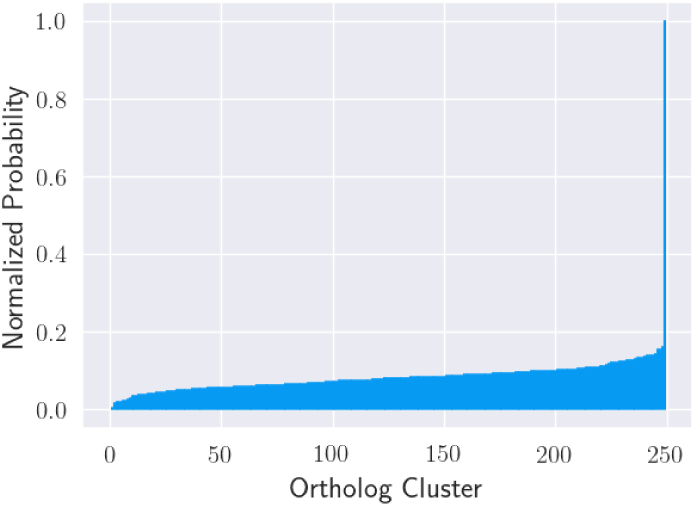
Distribution for valid clusters.

**Figure 3:**
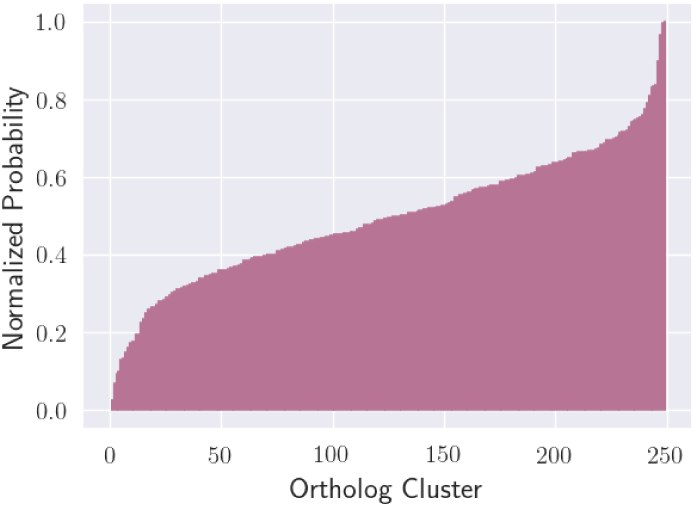
Distribution for outlier genes.

**Figure 4:**
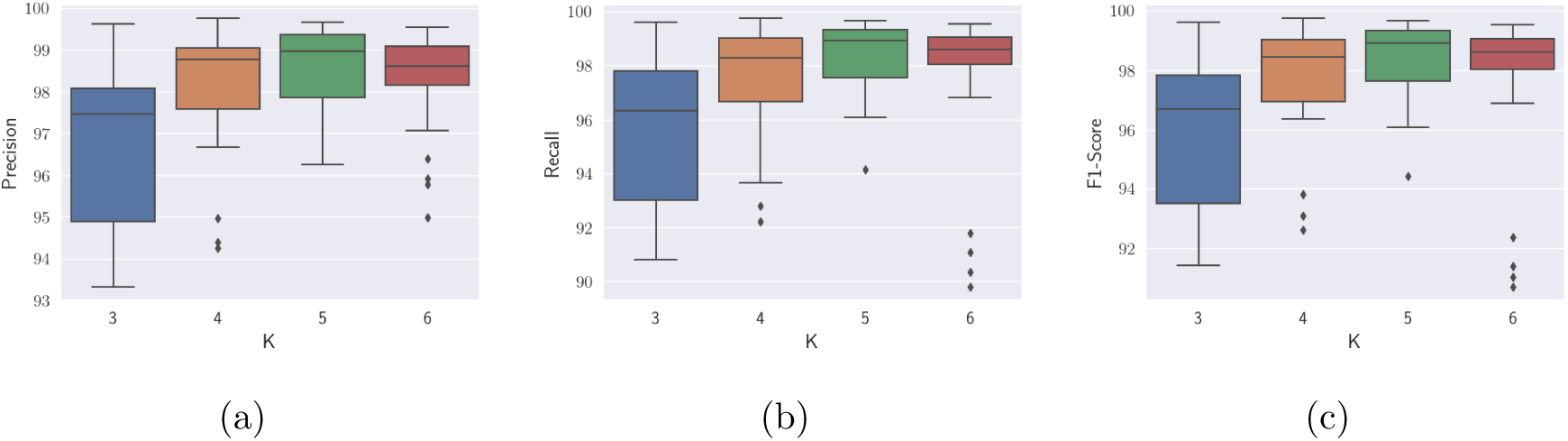
Effect of varying values of *k* on performance metrics (precision, recall and *F*_1_ score) on the KEGG dataset.

### 2.5 NORTH Software Availability

We implemented the Naive Bayes algorithm from scratch in python [52] without using the widely used Scikit-Learn [53] library as Scikit-Learn does not scale well to the amount of data we analyzed in this study. A server-side application using Flask[54] has been developed to aid the researchers for discovering the probable orthologs of a newly sequenced gene. We have also developed cross-platform, native desktop applications using Electron JS framework [55]. The server application as well as the desktop application can be accessed from the project website (https://nibtehaz.github.io/NORTH/).

## 3 Results

### 3.1 NORTH outperforms the frequently used methods on OrthoBench

One of the major complications in working with ortholog clustering is the lack of a standardized way to compare various methods since most authors have used different databases. Fortunately, OrthoBench [13] presents us a benchmark to compare various ortholog clustering methods. OrthoBench operates on a database of specially curated protein families of Metazoan organisms. It comprises a total of 1695 genes, distributed into 70 ortholog clusters. Particular attention was provided in selection of these clusters as 40 of the clusters present challenging biological scenarios. The remaining 30 clusters were randomly selected to eliminate any possible biases. OrthoBench originally tested five popular methods, but later OrthoFinder [22] was proven to outperform all of them.

NORTH was evaluated on this benchmark under a cross-validation scheme. We performed a stratified 5-fold cross-validation [56], such that circularly 80% of the data gets used as training data and the remaining 20% data is kept as test data. Selecting the stratified configuration prevented the data from omitting the small clusters. The results from cross-validation are remarkable, as NORTH overshadows all the six other methods. Most notably, it significantly improves upon the previous best performing method OrthoFinder, by achieving about 11.88% higher *F*_1_-score. Moreover, NORTH yielded an impressive trade-off between precision and recall, whereas methods like OMA, despite being the most precise, has the poorest recall.

### 3.2 NORTH manages to overcome the adversities in orthology inference

The most important consideration of the OrthoBench benchmark is exploring the effect of different challenges prevailing in ortholog clustering. OrthoBench benchmark studied at depth the impact of issues like bigger family sizes, faster rate of evolution, lower quality of sequence alignment and other challenging scenarios [13]. It was observed that all these conditions terribly affects the performance of ortholog clustering, resulting in increased numbers of false positives and false negatives, which were termed as *erroneously assigned genes* and *missing genes* respectively. The impact of these scenarios are summarized in Fig. 6. It is indisputably conspicuous that all the methods suffer miserably from these adverse conditions by incurring high levels of FPs and FNs, except for NORTH which is notably resilient against these adversities (please see Appendix B for more details).

**Figure 5:**
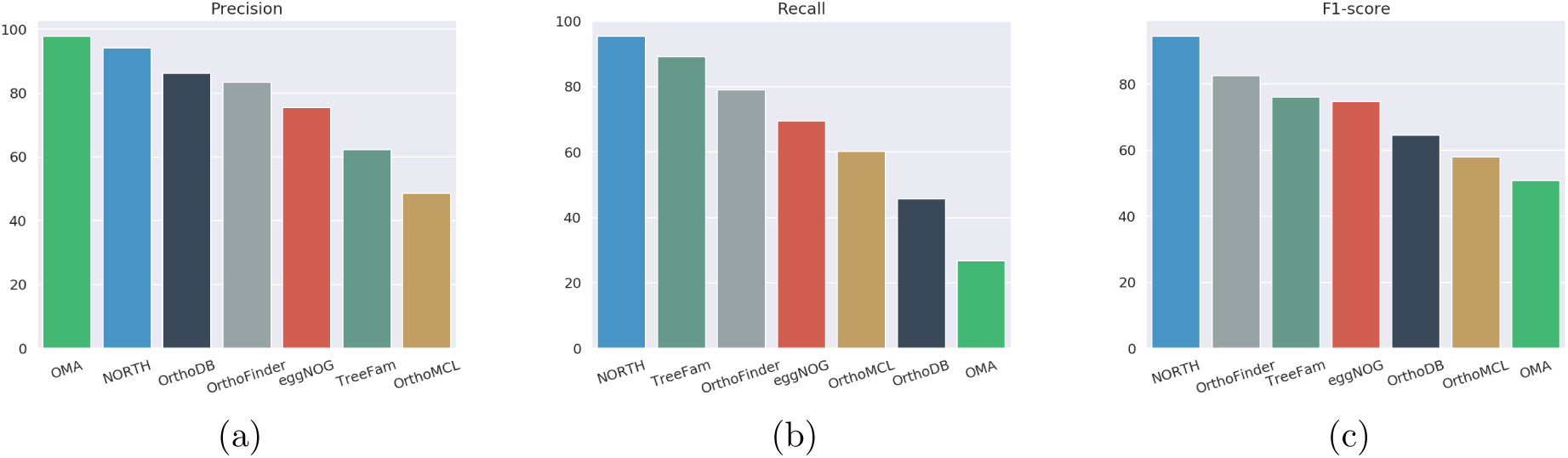
Performance on the OrthoBench benchmark. Figures (a) - (c) present the precision, recall and *F*_1_-score, respectively. From 5c, it is evident that NORTH is the best performing algorithm, having the highest *F*_1_-Score. The results are collected from the OrthoFinder paper [22].

**Figure 6:**
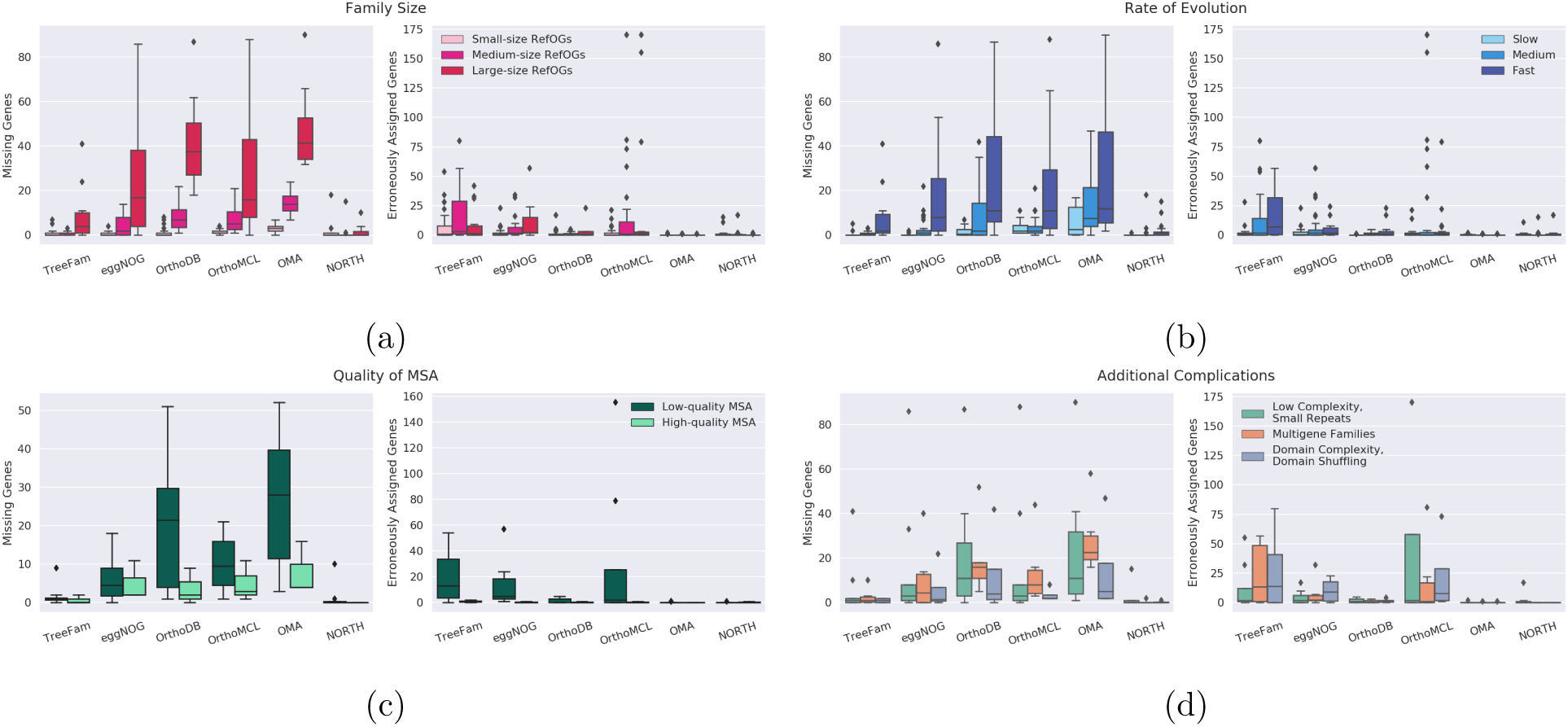
Performance evaluation under various Challenging scenarios in ortholog clustering. Figures (a) - (d) demonstrate the numbers of missing genes and erroneously assigned genes by various methods under different problematic circumstances.

### 3.3 NORTH is tested on diversified set of organisms

We have thoroughly tested NORTH on a vast repertoire of real-world orthologous cluster data from diverse organisms. We used the KEGG [57, 58] database for the most diverse collection of orthologous genes, which incorporates the molecular function of the genes as well through manual curation [59]. However, a challenge in analyzing the KEGG dataset is that the clusters therein are quite sparse. These clusters are highly imbalanced, with several of them having a few thousands of genes, while most of them have only a handful of genes (see Appendix C).

The KEGG database is also conveniently divided into several partitions, namely, Plants, Fungi, Protists, Animals, Bacteria, and Archaea, allowing us to test NORTH on all these different families of organisms. For all these different families, we take the biggest 10 clusters and gradually consider more clusters, i.e., 50, 100, 150, 200 and 250 biggest clusters, after which the number of genes in the clusters declines drastically. However, for Protists we considered only the biggest 10, 20, 30 and 40 clusters as its clusters are problematically small.

The results on various organisms are summarized in Fig. 7. Here, we present the values of *F*_1_-score for different families of organisms with different numbers of clusters under consideration. NORTH achieves impressively high *F*_1_-scores across various families, indicating a balance between precision and recall as well.

**Figure 7:**
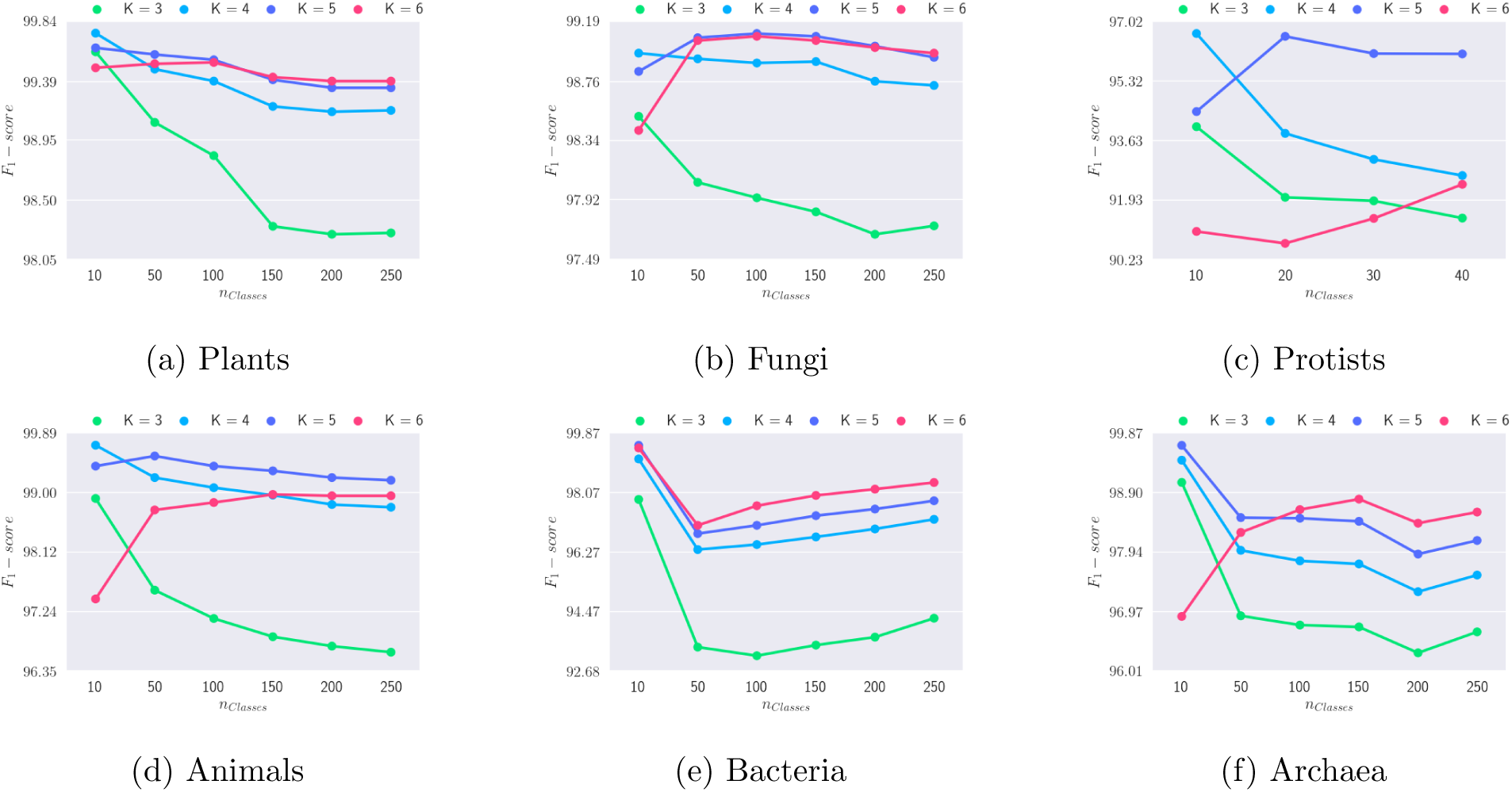
Performance of NORTH on various datasets in the KEGG database. We show results on the biggest 10, 50, 100, 150, 200 and 250 clusters (except for Protists). We reprot the *F*_1_ score for varying numbers of ortholog clusters with different values of *k* (3 ∼ 6) as it is a balanced representation of both precision and recall.

### 3.4 NORTH is consistent even when distant organisms are considered

Orthology prediction becomes excessively complicated when distantly related organisms are involved [24]. However, another remarkable property of NORTH is its capability of maintaining a high level of accuracy even when distantly related organisms are under consideration. To validate so, we performed experiments in two levels. First, we compiled all the eukaryotic genes and the prokaryotic genes and conducted experiments on these broader set of data in the similar manner as described in Sec. 3.3. Furthermore, we merged these two supersets as well and constructed a database of all the different types of organisms. Experimental outcomes (Fig. 8) reveal that even when considering such a voluminous spectrum of organisms, NORTH performs extremely well, suggested by the admiring values of *F*_1_-score.

**Figure 8:**
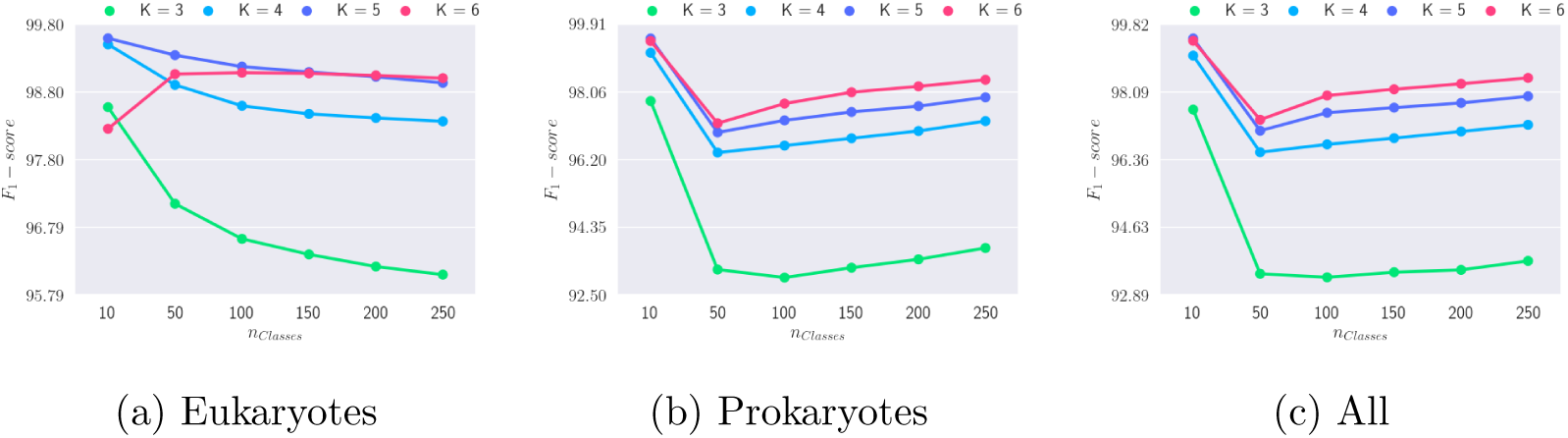
Performance of NORTH on distantly related organisms. We clustered all the eukaryotic (8a) and prokaryotic (8b) genes, both individually and jointly (8c).

### 3.5 NORTH is immune to data imbalance

One of the difficulties in analyzing biological datasets is the data imbalance [60]. Although machine learning based techniques are likely to fail to give robust predictions in the presence of data imbalance [61], our extensive experimental study establishes that NORTH is robust enough to handle data imbalance. When we consider the biggest 250 clusters, NORTH obtainspPrecision = 98.48%, recall = 98.43% and *F*_1_ score=98.44%, which is remarkable considering the intensity of data imbalance. It may be perceived that NORTH overfits the biggest clusters and thus obtains a high average accuracy. Figure 9, however, reveals that although only a few clusters have around 8, 000-24, 000 genes (majority with less than 4, 000), the performance metrics are above 90% for all but a few clusters. Therefore, NORTH is able to predict genes from even the smallest clusters accurately.

**Figure 9:**
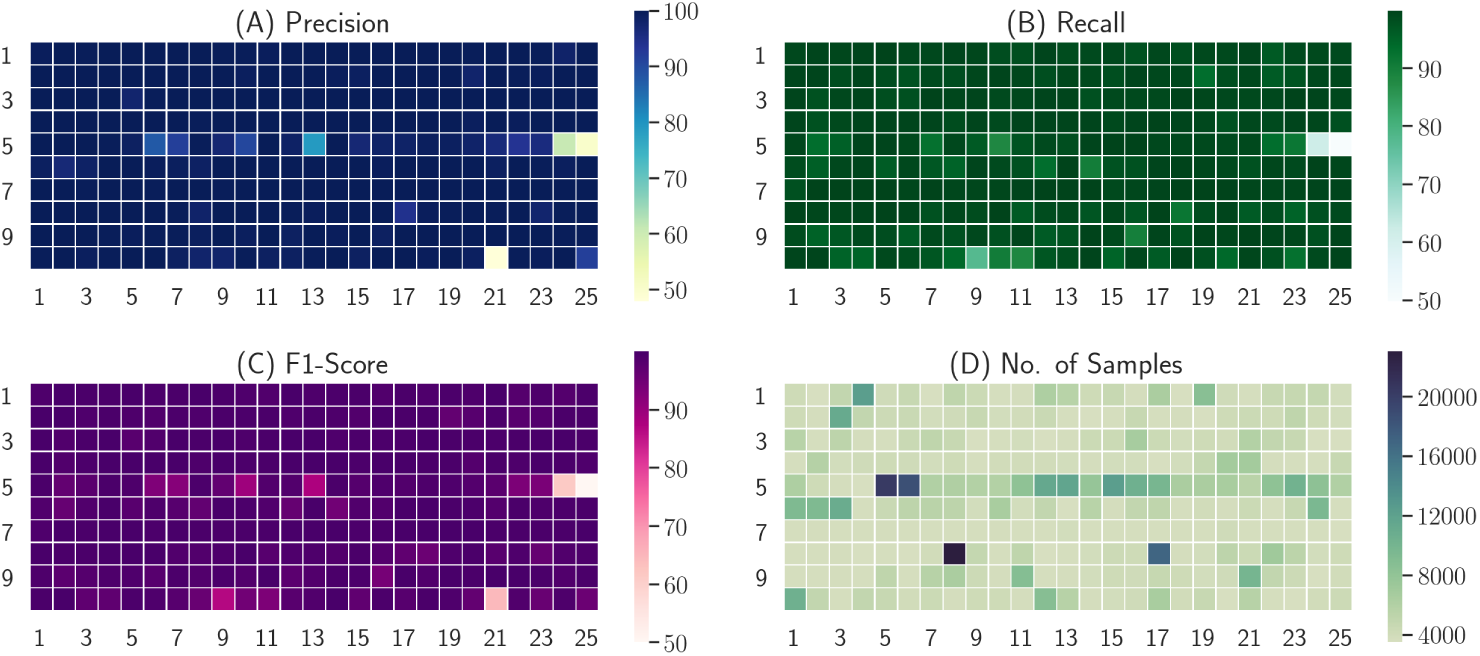
Performance of NORTH on highly imbalanced data. We show results on the largest 250 clusters comprising genes from all the six major groups of life. Here, the individual squares in the heatmaps correspond to different ortholog clusters.

### 3.6 NORTH is robust against outlier genes

To assess the effectiveness of NORTH in identifying genes that do not belong to the predefined set of clusters, we consider the biggest 250 clusters on all the organisms and train the Naive Bayes model. To introduce outliers, we consider all the remaining clusters with at least 3,000 genes. In this way, we obtain 12,55,877 member genes in the 250 clusters under consideration, and 6,48,938 genes from the clusters not within the 250 predefined ones. Consequently, features are computed and a Random Forest classifier comprising 10 trees is trained. The results of a stratified 10-fold cross validation are presented in Table 1. It is evident that NORTH has achieved a very high *F*_1_-score (98.67%), maintaining a good balance between precision and recall.

**Table 1:** Performance evaluation of the outlier detection. We effectively classify whether a gene is correctly assigned to a predefined cluster, or is an outlier.

| Class | Precision (%) | Recall (%) | $F_1$ score (%) |
| --- | --- | --- | --- |
| Outlier | 98.818 | 98.5365 | 98.6713 |
| Valid Clustering | 99.250 | 99.390 | 99.319 |
| Average | 99.103 | 99.099 | 98.6713 |

### 3.7 NORTH is scalable

In order to assess the scalability of NORTH, we ran different configurations of NORTH having different numbers of sub-processes (1 ∼ 8). For each configuration, a server process was dedicated to run an individual sub-process. All the models were trained on the biggest 250 eukaryote ortholog clusters from KEGG database. Next, 10,000 eukaryotic genes were selected randomly and the prediction time of different NORTH variants were computed. The average prediction time reduced from 31 ms to 14.5 ms when we divided the computation to 4 sub-processes instead of running it as a whole (see Fig. 10). However, as we divide the computation, additional overheads emerge. It was seen that for this volume of data, partitioning it further worsens the performance as the reduction of computation is outweighed by the overheads. However, it is intuitive that a higher number of sub-processes will be more effective for larger numbers of clusters. The server initialization time, which involves the loading of the models, is not considered here.

**Figure 10:**
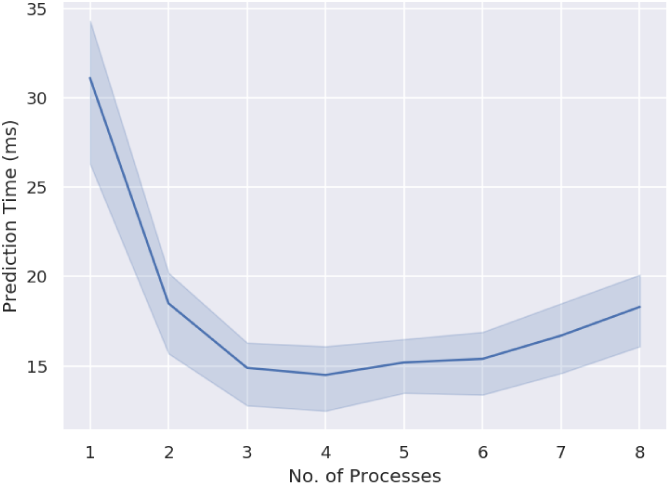
Scalability of NORTH.

## 4 Discussion and conclusions

In this work, we have made an attempt at developing a highly accurate and automatic machine learning based pipeline to cluster orthologous genes together. Orthology prediction is sufficiently complex and hence existing methods cannot fully comprehend or predict them. Thus the goal here should be the creation of an appropriate model; the model should account for as much genome data as possible, and continue improving by modifying itself to incorporate new data and scientific findings. This view emphasizes the importance of machine learning in orthology identification. ML-based algorithms allow us to analyze a plethora of data, which is not possible to analyze manually. Moreover, these algorithms try to adaptively select the best set of parameters, and thus remove human biases and the necessity of manual intervention while setting various parameters. Therefore, machine learning algorithms are more promising for developing algorithms in comparative genomics due to the availability of a vast repertoire of gene sequences. However, the existing ML-based techniques mostly depend on sophisticated feature engineering, are prone to precision-recall trade-off and constrained to small numbers of organisms – making them impractical for handling large amount of complex biological data.

NORTH involves a novel combination of tools from natural language processing and machine learning with the biological basis of orthologs, and offers not only high accuracy which is otherwise quite hard to achieve without sophisticated manual intervention, but scalability that enables successful analyses of millions of genes on a wide range of distantly related organisms without having to rely on supercomputers. NORTH works satisfactorily even in a core i3 CPU with 4 GB of RAM.

BLAST algorithm demands high processing memory, restricting the methods to limited numbers of sequences. Scalability and reduced running time of NORTH can be attributed to replacing the expensive BLAST search and subsequent graph based algorithms, and our highly efficient and customized in-house implementation of the multinomial Naive Bayes algorithm.

Results on the OrthoBench dataset clearly shows its superiority over the best alternate methods and demonstrate its robustness against various challenging scenarios which badly affect other methods. NORTH was able to analyze over a million of genes in KEGG database and achieved excellent accuracy. Although, due to the lack of sufficient data in individual clusters, we have considered the largest 250 clusters, we have shown that NORTH can identify genes that do not belong to these 250 predefined clusters with high precision and recall. We expect the number of available genomes to expand rapidly and to encompass new genes in improving our system. NORTH is currently not suitable for pairwise orthology prediction. Although a clustering algorithm can also infer the pairwise relationship, the one-to-one, one-to-many, and many-to-many variants of orthologs need to be considered carefully as they may raise false negatives or false positives. We left this as a future work.

On an ending note, we have demonstrated that an appropriate bag-of-words model with a Naive Bayes classifier can successfully cluster the orthologous genes together. This study is the first of its kind, and we believe this represents a significant advance in identification of orthologous genes. NORTH will evolve with the availability of new data, and in response to scientific findings and systamatists’ feedback – laying a firm, broad foundation for fully automated, highly accurate and scalable orthology identification.

## Appendix A Overview of the scalable Naive Bayes implementation

We use a multinomial Naive Bayes model to cluster the orthologs. The problem is to classify a gene sequence *g* to one of the *m* ortholog clusters {*C*_1_, *C*_2_, *C*_3_, …, *C*_*m*_}. The input to the Naive Bayes model is a vector *x* = [*x*_1_, *x*_2_, *x*_3_, …, *x*_*n*_] that contains the *k*-mer frequencies of a given gene *g.* Here, *x*_*i*_ represents the frequency of the *i*-th *k*-mer and *n* is the total number of *k*-mers. The naive Bayes model predicts the probability of a gene sequence with *k*-mer frequency *x*, to be a member of cluster *C*_*k*_ as *p*(*C*_*k*_|*x*),

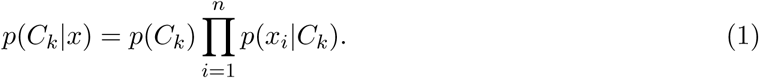

Here, *p*(*C*_*k*_) is the prior probabilty of cluster *C*_*k*_ and *p*(*x*_*i*_|*C*_*k*_) is the likelihood of the *i*-th *k*-mer. These terms are learnt during the training phase using the following formulae.

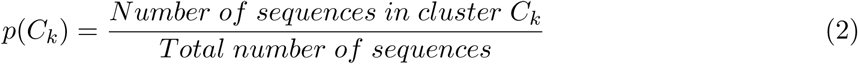

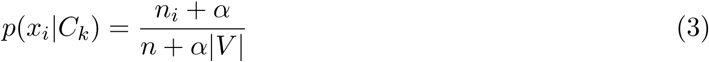

Here, *n*_*i*_ = number of occurrences of *x*_*i*_ in *C*_*K*_, *α* = smoothing factor (we used *α* = 1, which corresponds to Laplace smoothing [47]), *n* = total number of *k*-mers in cluster *C*_*k*_, |*V* | = size of the vocabulary (number of unique *k*-mers in the entire data).

During the testing phase, i.e. obtaining predicted clusters for an unknown gene *g*′ with *k*-mer frequency *x*′, we compute the probabilities of *g*′ being a member of the individual clusters *p*(*C*_*k*_|*x*′) for *k* = 1 … *m*. The cluster with the highest probability is the most likely solution *Ĉ*.

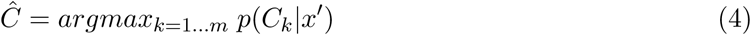

The training phase of Naive Bayes algorithm has a time complexity of *O*(*G* × *n*) and memory requirement of *O*(*n* × *m*), where *G* = number of genes in training data, *n* = number of *k*-mers and *m* = number of clusters. The time complexity of making a prediction in testing phase is *O*(*n* × *m*). Therefore, the algorithm becomes computationally quite expensive for large numbers of clusters with diversified *k*-mers. However, during both training and testing phases, the computations for the individual clusters are independent of each other and thus can be performed in parallel.

Thus, we divide all our computations into *p* child processes or jobs. A child process or job *p*_*i*_ is responsible for the computations of the clusters 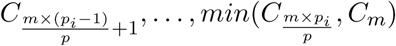, for 1 ≤ *p*_*i*_ ≤ *p*. Hence, during both training and testing phases, the computations are mapped to various parallel jobs and finally combined together. This is similar to the *map-reduce framework*^1^. Individual jobs can be run in parallel as multiple processes, multiple threads or even as multiple systems in a computer cluster. It reduces the memory requirement from *O*(*n* × *m*) to 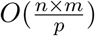 for the individual jobs. Moreover, individual jobs can be computed in a shorter amount of time 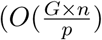 for training and 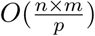 for testing). An schematic of the scalable implementation of NORTH is presented in Fig. 11.

**Figure 11:**
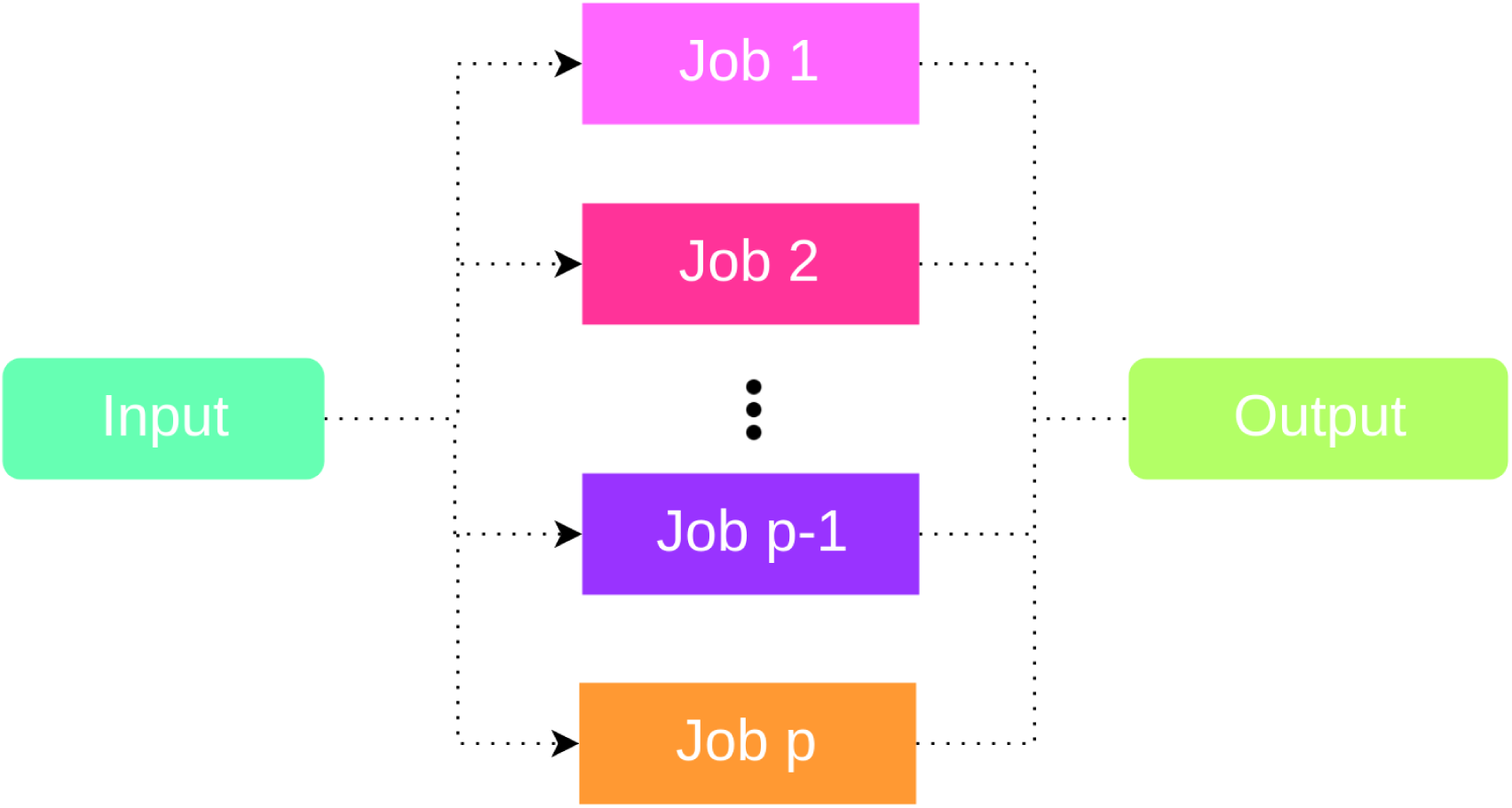
Scalable naive Bayes implementation.

In a limited computing resource setting, the jobs can be done sequentially, which keeps the memory requirement manageable at roughly the same computation time (considering some over-heads). In the presence of sufficient computational resources, the jobs can be done in parallel as described above, enabling us to analyze large numbers of ortholog clusters.

## Appendix B OrthoBench

OrthoBench has been comprehensively explained in the original paper [13]. The data for OrthoBench was downloaded from the project webpage http://eggnog.embl.de/orthobench/. In [13], the authors particularly investigated the effect of the following challenging criteria.

- Family size: the authors divided 70 orthologous clusters into 3 categories based on the family size, (i) small size (less than 14 members), (ii) medium size (more than 14 but less than 40 members) and (iii) large size (more than 40 members). It was demonstrated that as the family size increases, the performance of various algorithms declines.
- Rate of evolution: the benchmarking clusters were categorized into slow-, medium-, and fast-evolving based on their MeanID score (described as the FamID in [62]) as follows: (i) slow-evolving (MeanID > 0.7), (ii) medium-evolving (0.5 < MeanID < 0.7), and (iii) fast-evolving (MeanID < 0.5).
- Quality of MSA: the authors considered 11 clusters based on their alignment quality. 8 of them were of low quality and the remaining 3 were of high quality.
- Domain architecture complexity: They also analyzed the domain complexity of the protein families. However, due to the lack of sufficient information in the original paper, supplementary materials or the project website, we could not reproduce and analyze this model condition.
- Miscellaneous: the authors also pointed out some challenging scenarios such as, low complexity and small repeats, multigene families, domain complexity and domain shuffling.

The numbers of FPs (erroneously classified genes) and FNs (missing genes) of the five methods were obtained from the supplementary materials of [13]. We were not able to include OrthoFinder in this comparison as it was not assessed under these challenging scenarios.

## Appendix C Results on the KEGG database

Among the ortholog databases, the KEGG [57, 58] database offers the most diverse collection of orthologous genes. These genes are arranged in a substantially large number of orthologous clusters. When most other databases available in the literature solely depend on sequence similarity measures [59], KEGG database incorporates the molecular function of the genes as well. This manual curation makes the KEGG dataset superior compared to other well-studied databases based on a BLAST search based protocol [59]. However, a challenge in analyzing the KEGG dataset is that the clusters therein are quite sparse. In fact, these clusters are highly imbalanced, with several having a few thousands of genes while some other with only a handful (see Fig. 12). There are 7,895 clusters among which, the biggest cluster contains 23,012 genes, whereas the smallest ones contain single gene only. The mean, median and standard deviation of the gene distributions in various clusters are 691, 220 and 1,195 respectively, indicating the presence of acute imbalance. Thus, we handled a big challenge of training machine learning algorithm on imbalanced datasets [61].

**Figure 12:**
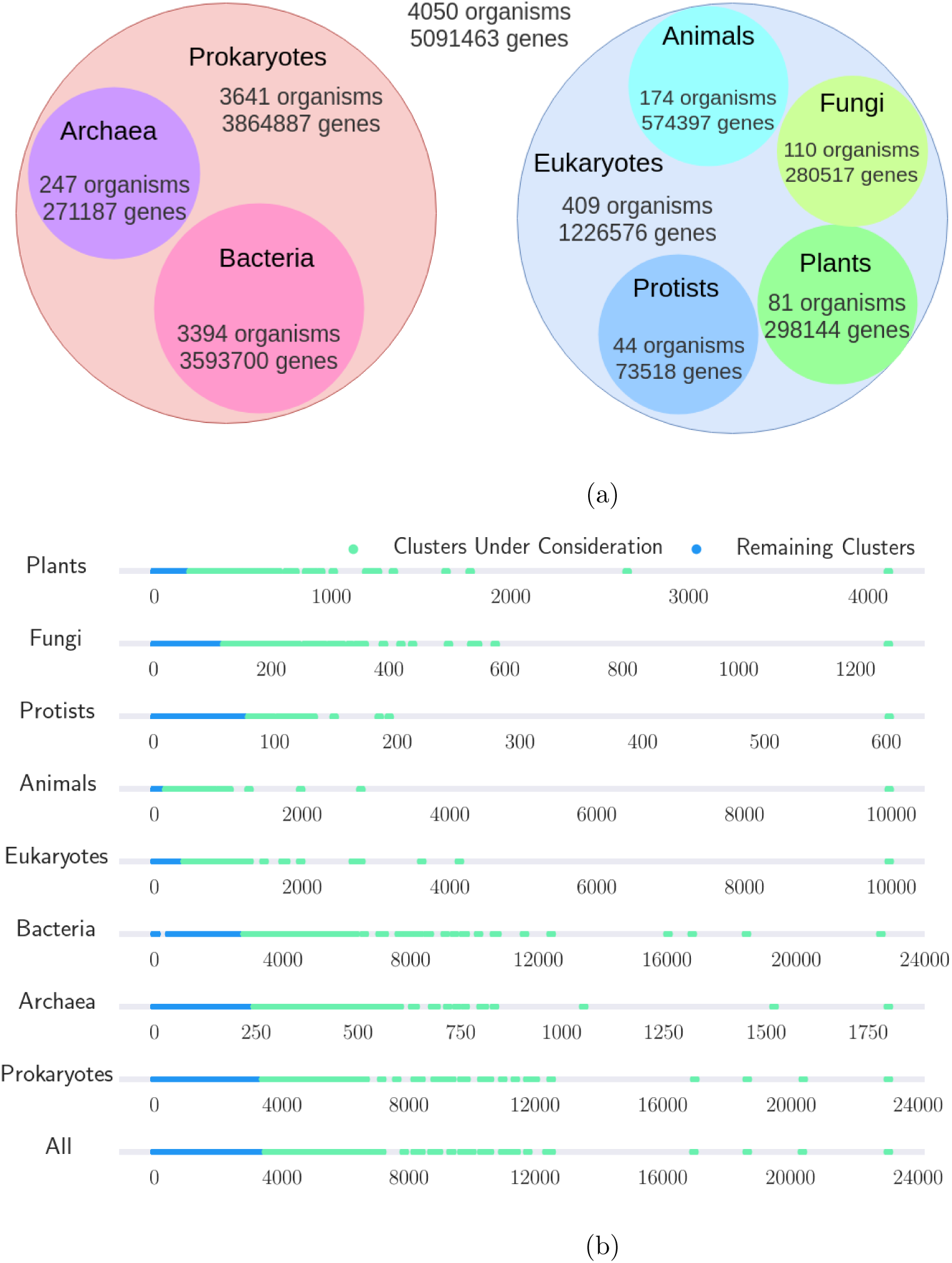
Overview of the KEGG dataset. (a) Distribution of organisms and genes in the nine sub-groups of life in KEGG database. (b) Distribution of gene population among different ortholog clusters, showing the number of genes in each of the clusters. We considered the biggest 250 clusters from each of the sub-groups (except for ‘Protists’ where we considered the largest 40).

The KEGG database is conveniently divided into several partitions, namely, Plants, Fungi, Protists, Animals, Eukaryotes, Bacteria, Archaea, Prokaryotes, and a partition containing all the organisms (refered to as ‘All’). This allowed us to test NORTH on different families of species and validate NORTH on diverse species families. We manually collected the ortholog tables from KEGG web browser [63], and then fetched the protein sequences from UniProt [64] using the UniProt REST API [65]. Unfortunately, a few genes listed in KEGG were not present in UniProt, which we had to exclude from our experiments. The overview of this dataset is depicted in Fig. 12a, showing the distributions of organisms and genes among various groups of life. In Fig. 12b, we have illustrated the data imbalance among the clusters. The gene population in the biggest clusters are scattered over large ranges of gene counts, and the majority of the remaining clusters contain only a handful of genes. This acute data imbalance prevented us from studying all the clusters and limited our investigation to only the biggest clusters.

The summary of the results of our experiments based on the KEGG database are presented here. More detailed experimental results with performance metrics (precision, recall and *F*_1_ score) for each of the individual orthologous clusters are provided in the supplementary material.

## Appendix D Independent testing

In order to perform an independent testing (performance of NORTH on the unseen data), we considered the largest 250 clusters and from each of the clusters, we took 10% of the total samples at random to form the independent testing set. This ensured adequate presence of genes from all the clusters in the test set. Using the rest of the data (i.e., training data in this context), we performed a stratified 10-fold cross validation. Next, we used the best performing model (based on the *F*_1_-score) to predict the clusters of the genes from the test set. We also performed an ensemble (majority voting) of all the ten models, resulted from the stratified 10-fold cross validation, and observed the performance on the test set. The results are presented in Table 11. These results indicate the generality of NORTH, as the performance of the best model is similar to the ensemble of all the trained models, and the performance metrics are around 98%.

**Table 2:** Performance on the Plants dataset. We vary the numbers of clusters from 10 to 250 and show Precision, Recall and *F*_1_-score for various values of *k* (3 ∼ 6). Best results under each model condition are shown in bold.

| $n_{clusters}$ | $k = 3$ | | | $k = 4$ | | | $k = 5$ | | | $k = 6$ | | | Total Genes |
| --- | --- | --- | --- | --- | --- | --- | --- | --- | --- | --- | --- | --- | --- |
| | Pr | Re | $F_1$ | Pr | Re | $F_1$ | Pr | Re | $F_1$ | Pr | Re | $F_1$ | |
| 10 | 99.61 | 99.61 | 99.61 | **99.75** | **99.75** | **99.75** | 99.65 | 99.64 | 99.64 | 99.5 | 99.49 | 99.49 | 17351 |
| 50 | 99.18 | 99.04 | 99.08 | 99.49 | 99.47 | 99.48 | **99.59** | **99.59** | **99.59** | 99.53 | 99.52 | 99.52 | 39534 |
| 100 | 98.98 | 98.78 | 98.83 | 99.41 | 99.39 | 99.39 | **99.55** | **99.55** | **99.55** | 99.53 | 99.53 | 99.53 | 56217 |
| 150 | 98.6 | 98.2 | 98.3 | 99.23 | 99.2 | 99.2 | 99.41 | 99.4 | 99.4 | **99.43** | **99.42** | **99.42** | 69176 |
| 200 | 98.56 | 98.12 | 98.24 | 99.19 | 99.15 | 99.16 | 99.35 | 99.35 | 99.34 | **99.39** | **99.39** | **99.39** | 80764 |
| 250 | 98.59 | 98.12 | 98.25 | 99.2 | 99.16 | 99.17 | 99.35 | 99.34 | 99.34 | **99.39** | **99.39** | **99.39** | 89690 |

**Table 3:** Performance on the Fungi dataset. We vary the numbers of clusters from 10 to 250 and show Precision, Recall and *F*_1_-score for various values of *k* (3 ∼ 6). Best results under each model condition are shown in bold.

| $n_{clusters}$ | $k = 3$ | | | $k = 4$ | | | $k = 5$ | | | $k = 6$ | | | Total Genes |
| --- | --- | --- | --- | --- | --- | --- | --- | --- | --- | --- | --- | --- | --- |
| | Pr | Re | $F_1$ | Pr | Re | $F_1$ | Pr | Re | $F_1$ | Pr | Re | $F_1$ | |
| 10 | 98.53 | 98.51 | 98.51 | **98.96** | **98.96** | **98.96** | 98.86 | 98.82 | 98.83 | 98.57 | 98.37 | 98.41 | 5759 |
| 50 | 98.18 | 98.0 | 98.04 | 98.96 | 98.91 | 98.92 | 99.09 | **99.07** | **99.07** | **99.24** | 98.97 | 99.05 | 16381 |
| 100 | 98.13 | 97.87 | 97.93 | 98.92 | 98.88 | 98.89 | **99.11** | **99.09** | **99.1** | 99.1 | 99.07 | 99.08 | 26268 |
| 150 | 98.09 | 97.75 | 97.83 | 98.94 | 98.89 | 98.9 | **99.09** | **99.08** | **99.08** | 99.06 | 99.05 | 99.05 | 33642 |
| 200 | 97.99 | 97.57 | 97.67 | 98.82 | 98.75 | 98.76 | **99.02** | **99.0** | **99.01** | 99.01 | **99.0** | 99.0 | 40810 |
| 250 | 98.06 | 97.61 | 97.73 | 98.8 | 98.71 | 98.73 | 98.95 | 98.93 | 98.93 | **98.97** | **98.96** | **98.96** | 47952 |

**Table 4:** Performance on the Protists dataset. We vary the numbers of clusters from 10 to 40 and show Precision, Recall and *F*_1_-score for various values of *k* (3 ∼ 6). Best results under each model condition are shown in bold.

| $n_{clusters}$ | $k = 3$ | | | $k = 4$ | | | $k = 5$ | | | $k = 6$ | | | Total Genes |
| --- | --- | --- | --- | --- | --- | --- | --- | --- | --- | --- | --- | --- | --- |
| | Pr | Re | $F_1$ | Pr | Re | $F_1$ | Pr | Re | $F_1$ | Pr | Re | $F_1$ | |
| 10 | 95.5 | 93.87 | 94.01 | **97.02** | **96.64** | **96.66** | 96.6 | 94.14 | 94.44 | 95.92 | 90.35 | 91.03 | 1876 |
| 20 | 94.24 | 91.43 | 92.0 | 94.97 | 93.64 | 93.82 | **96.96** | **96.5** | **96.58** | 96.38 | 89.79 | 90.69 | 2939 |
| 30 | 93.77 | 91.38 | 91.9 | 94.38 | 92.81 | 93.08 | **96.26** | **96.08** | **96.09** | 94.98 | 91.07 | 91.4 | 3851 |
| 40 | 93.32 | 90.81 | 91.41 | 94.25 | 92.2 | 92.62 | **96.28** | **96.08** | **96.08** | 95.78 | 91.78 | 92.37 | 4669 |

**Table 5:**
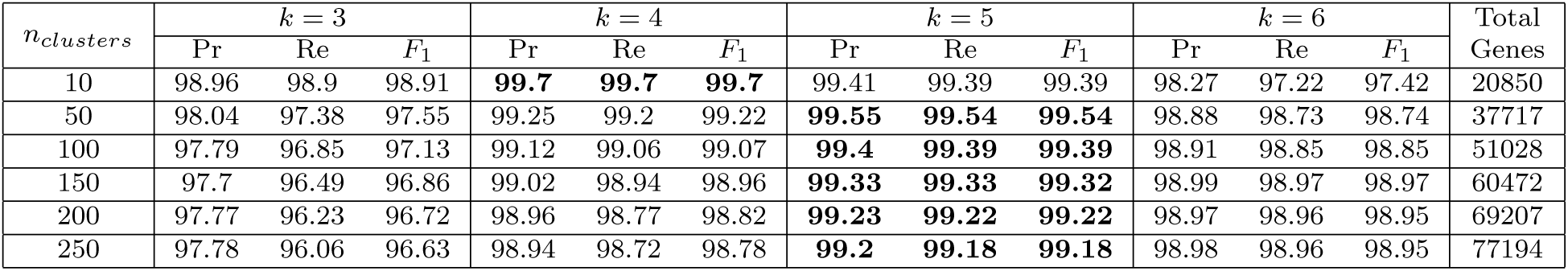
Performance on the Animals dataset. We vary the numbers of clusters from 10 to 250 and show Precision, Recall and *F*_1_-score for various values of *k* (3 ∼ 6). Best results under each model condition are shown in bold.

**Table 6:** Performance on the Eukaryotes dataset. We vary the numbers of clusters from 10 to 250 and show Precision, Recall and *F*_1_-score for various values of *k* (3 ∼ 6). Best results under each model condition are shown in bold.

| $n_{clusters}$ | $k = 3$ | | | $k = 4$ | | | $k = 5$ | | | $k = 6$ | | | Total Genes |
| --- | --- | --- | --- | --- | --- | --- | --- | --- | --- | --- | --- | --- | --- |
| | Pr | Re | $F_1$ | Pr | Re | $F_1$ | Pr | Re | $F_1$ | Pr | Re | $F_1$ | |
| 10 | 98.66 | 98.54 | 98.57 | 99.51 | 99.5 | 99.5 | **99.6** | **99.58** | **99.59** | 98.6 | 98.18 | 98.25 | 31787 |
| 50 | 97.48 | 97.05 | 97.14 | 98.94 | 98.88 | 98.9 | **99.35** | **99.34** | **99.34** | 99.1 | 99.05 | 99.06 | 70791 |
| 100 | 97.57 | 96.13 | 96.62 | 98.85 | 98.48 | 98.59 | **99.19** | **99.17** | **99.17** | 99.09 | 99.08 | 99.08 | 103229 |
| 150 | 97.48 | 95.8 | 96.39 | 98.8 | 98.32 | 98.47 | **99.11** | **99.08** | **99.09** | 99.08 | 99.07 | 99.07 | 129805 |
| 200 | 97.42 | 95.56 | 96.21 | 98.74 | 98.27 | 98.41 | **99.05** | 99.01 | 99.02 | **99.05** | **99.04** | **99.04** | 153368 |
| 250 | 97.36 | 95.42 | 96.09 | 98.7 | 98.2 | 98.36 | 98.98 | 98.92 | 98.93 | **99.01** | **99.0** | **99.0** | 175088 |

**Table 7:** Performance on the Bacteria dataset. We vary the numbers of clusters from 10 to 250 and show Precision, Recall and *F*_1_-score for various values of *k* (3 ∼ 6). Best results under each model condition are shown in bold.

| $n_{clusters}$ | $k = 3$ | | | $k = 4$ | | | $k = 5$ | | | $k = 6$ | | | Total Genes |
| --- | --- | --- | --- | --- | --- | --- | --- | --- | --- | --- | --- | --- | --- |
| | Pr | Re | $F_1$ | Pr | Re | $F_1$ | Pr | Re | $F_1$ | Pr | Re | $F_1$ | |
| 10 | 97.93 | 97.82 | 97.85 | 99.1 | 99.06 | 99.07 | **99.49** | **99.48** | **99.48** | 99.4 | 99.4 | 99.4 | 138423 |
| 50 | 94.23 | 93.07 | 93.4 | 96.66 | 96.24 | 96.34 | 96.87 | 96.81 | 96.82 | **97.07** | **97.08** | **97.07** | 395363 |
| 100 | 94.42 | 92.55 | 93.14 | 97.04 | 96.27 | 96.49 | 97.27 | 97.02 | 97.07 | **97.68** | **97.66** | **97.66** | 602179 |
| 150 | 94.84 | 92.77 | 93.46 | 97.36 | 96.44 | 96.72 | 97.59 | 97.28 | 97.36 | **98.0** | **97.97** | **97.97** | 773812 |
| 200 | 95.05 | 93.02 | 93.7 | 97.59 | 96.67 | 96.96 | 97.82 | 97.46 | 97.56 | **98.19** | **98.16** | **98.16** | 924677 |
| 250 | 95.52 | 93.64 | 94.27 | 97.83 | 96.98 | 97.25 | 98.06 | 97.72 | 97.81 | **98.39** | **98.35** | **98.36** | 1065520 |

**Table 8:** Performance on the Archaea dataset. We vary the numbers of clusters from 10 to 250 and show Precision, Recall and *F*_1_-score for various values of *k* (3 ∼ 6). Best results under each model condition are shown in bold.

| $n_{clusters}$ | $k = 3$ | | | $k = 4$ | | | $k = 5$ | | | $k = 6$ | | | Total Genes |
| --- | --- | --- | --- | --- | --- | --- | --- | --- | --- | --- | --- | --- | --- |
| | Pr | Re | $F_1$ | Pr | Re | $F_1$ | Pr | Re | $F_1$ | Pr | Re | $F_1$ | |
| 10 | 99.07 | 99.06 | 99.06 | 99.43 | 99.42 | 99.42 | **99.66** | **99.66** | **99.66** | 97.41 | 96.8 | 96.89 | 9689 |
| 50 | 97.19 | 96.81 | 96.9 | 98.05 | 97.94 | 97.96 | **98.54** | **98.49** | **98.49** | 98.33 | 98.24 | 98.25 | 29957 |
| 100 | 97.21 | 96.59 | 96.75 | 98.0 | 97.76 | 97.79 | 98.58 | 98.47 | 98.48 | **98.63** | **98.62** | **98.62** | 47174 |
| 150 | 97.23 | 96.53 | 96.72 | 98.04 | 97.67 | 97.74 | 98.6 | 98.4 | 98.43 | **98.8** | **98.8** | **98.79** | 61533 |
| 200 | 96.87 | 96.08 | 96.3 | 97.67 | 97.19 | 97.29 | 98.14 | 97.84 | 97.9 | **98.41** | **98.4** | **98.4** | 74649 |
| 250 | 97.19 | 96.42 | 96.64 | 97.95 | 97.44 | 97.56 | 98.38 | 98.05 | 98.12 | **98.59** | **98.58** | **98.58** | 90708 |

**Table 9:**
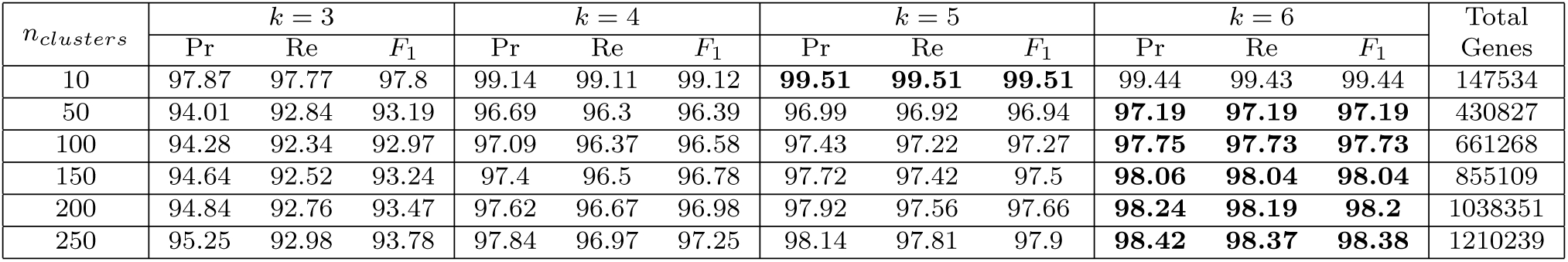
Performance on the Prokaryotes dataset. We vary the numbers of clusters from 10 to 250 and show Precision, Recall and *F*_1_-score for various values of *k* (3 ∼ 6). Best performance metrics under each model condition are shown in bold.

**Table 10:** Performance on the entire dataset (i.e., All dataset) comprising six major groups of life (Archaea, Bacteria, Animals, Fungi, Plants and Protists). We vary the numbers of clusters from 10 to 250 and show Precision, Recall and *F*_1_-score for various values of *k* (3 ∼ 6). Best results under each model condition are shown in bold.

| $n_{clusters}$ | $k = 3$ | | | $k = 4$ | | | $k = 5$ | | | $k = 6$ | | | Total Genes |
| --- | --- | --- | --- | --- | --- | --- | --- | --- | --- | --- | --- | --- | --- |
| | Pr | Re | $F_1$ | Pr | Re | $F_1$ | Pr | Re | $F_1$ | Pr | Re | $F_1$ | |
| 10 | 97.7 | 97.6 | 97.63 | 99.03 | 99.01 | 99.01 | **99.46** | **99.45** | **99.45** | 99.39 | 99.39 | 99.39 | 148591 |
| 50 | 94.24 | 93.08 | 93.43 | 96.83 | 96.45 | 96.54 | 97.14 | 97.08 | 97.09 | **97.37** | **97.37** | **97.37** | 442391 |
| 100 | 94.5 | 92.75 | 93.34 | 97.24 | 96.53 | 96.74 | 97.68 | 97.51 | 97.55 | **97.99** | **97.99** | **97.99** | 684238 |
| 150 | 94.79 | 92.78 | 93.47 | 97.5 | 96.62 | 96.9 | 97.87 | 97.61 | 97.68 | **98.17** | **98.15** | **98.15** | 888581 |
| 200 | 94.91 | 92.84 | 93.53 | 97.68 | 96.77 | 97.07 | 98.03 | 97.72 | 97.8 | **98.33** | **98.29** | **98.29** | 1076332 |
| 250 | 95.12 | 93.06 | 93.76 | 97.86 | 96.94 | 97.24 | 98.21 | 97.88 | 97.97 | **98.48** | **98.43** | **98.44** | 1255877 |

**Table 11:** Performance evaluation using an independent testing.

| Model | Precision | Recall | $F_1$ score |
| --- | --- | --- | --- |
| Best Model | 98.15 | 97.81 | 97.90 |
| Ensemble (majority voting) | 98.14 | 97.82 | 97.91 |

## Appendix E Declarations

### Supplementary Material

The supplementary material (SM.xls) presents the detailed results showing the performance metrics for each of the individual ortholog clusters under various model conditions.

### Data and code availability

The datasets analysed during the current study are available at KEGG (https://www.genome.jp/kegg/), and UniProt (https://www.uniprot.org/) databases. The NORTH software is freely available as open source code at https://github.com/nibtehaz/NORTH. A step-by-step, comprehensive tutorial for NORTH is available at http://nibtehaz.github.io/NORTH-app

### Author contributions

MSB conceived the study and NI helped design the study; NI, SA, BS, MSB and MSR developed the methods; NI implemented the methods; NI, SA, and BS conducted the experiments; NI, MSB and MSR analyzed and interpreted the results; MSB and MSR supervised the study; NI and MSB wrote the first draft and all the authors took part in finalizing the manuscript.

## Footnotes

1 https://ai.google/research/pubs/pub62

